# Profiles and clinical significance of immune cell infiltration in pancreatic adenocarcinoma

**DOI:** 10.1101/2020.03.30.017327

**Authors:** Jie Mei, Rui Xu, Dandan Xia, Xuejing Yang, Huiyu Wang, Chaoying Liu

## Abstract

**Background:** It has been well defined that tumor-infiltrating immune cells (TIICs) play critical roles in pancreatic cancer (PAAD) progression. The aim of this research was to comprehensively explore the composition of TIICs in PAAD and their potential clinical significance.

**Methods:** 178 samples from TCGA and 63 samples from GSE57495 dataset were enrolled into our study. ImmuCellAI was applied to calculate the infiltrating abundance of 24 immune cell types in PAAD and further survival analysis revealed the prognostic values of TIICs in PAAD. Moreover, Gene ontology (GO) enticement analysis of differentially expressed genes (DEGs) between low- and high-risk groups was performed as well.

**Results:** Different kinds of TIICs had distinct infiltrating features. Besides, Specific TIICs subsets had notable prognostic values in PAAD. We further established a 6-TIICs signature to assess the prognosis of PAAD patients. Kaplan-Meier and Cox regression analyses both suggested the significant prognostic value of the signature in PAAD. We next extracted 1,334 DEGs based on the risk model, and the hub modules in the protein-protein interaction (PPI) network of DEGs were involved in regulating immune-related biological processes.

**Conclusions:** Overall, the current study illuminated the immune cells infiltrating landscape in PAAD and developed a TIICs-dependent prognostic signature, which could be used as an effective prognostic classifier for PAAD patients.

## 1. Introduction

Pancreatic cancer (PAAD) is one of the most fatal cancerous disease worldwide, which is characterized with dreadful aggressiveness and poor prognosis. The early diagnosis of PAAD is difficult due to the obscure symptoms, and its morbidity and mortality have been increasing significantly in recent years [1]. According to the newest statistic data published by American Cancer Society, there will be about 57,600 new PAAD cases and nearly 50,000 cancer-causing deaths in 2020 in the USA [1]. With the constant development of advanced therapeutic strategies, immunotherapy is becoming a novel promising hotspot in the field of PAAD treatment [2]. The tumor immune microenvironment, which containing extracellular matrix, fibroblasts, endothelial cells, and multiple immune cells, plays a critical role in determining response to immunotherapy [3]. Increasing evidence reveals that tumor progression is significantly affected by host immune response, which is represented by abundance of tumor-infiltrating immune cells (TIICs) [4, 5]. However, the profiles and clinical significance of TIICs in PAAD have not been well defined.

In the past decades, high-throughput sequencing technologies, including microarray and RNA sequencing (RNA-Seq), produce massive transcriptome data, which makes estimating the abundance of TIICs by gene expression data possible. Several classic algorithms, including CIBERSORT [6], xCell [7], TIMER[8], EPIC [9], and MCPcounter [10] have been established to calculate immune cells abundance based on transcriptome data of tumor samples. Recently, Miao *et al*. developed a highly accurate method named ImmuCellAI to estimate the infiltrating levels of immune cells from transcriptome data [11]. ImmuCellAI expand the scope of infiltrating assessment of more T cell subsets, such as regulatory cell (Treg), cytotoxic T cell (Tc), and exhausted T cell (Tex). Besides, compared with other methods, ImmuCellAI has the highest consistency with flow cytometry results for most immune cells.

In this research, based on the ImmuCellAI approach and PAAD transcriptome data from TCGA and GEO datasets, we conducted an in-depth analysis of the TIICs in PAAD samples. As a result, six kinds of immune cells, including nTreg, T helper 1 cell (Th1), Th17, dendritic cell (DC), CD4+ T cell, and CD8+ T cell, was developed as promising predictive signature for prognostic assessment for PAAD patients. Furthermore, we analyzed the risk-associated differentially expressed genes (DEGs), performed the functional enrichment analysis, and constructed the protein-protein interaction (PPI) network, provided a novel understand about the progression of PAAD from the insight of immune-related biological processes.

## 2. Materials and methods

### 2.1. Data acquisition

Normalized RNA-seq data and clinical information of PAAD samples were downloaded from UCSC Xena website (https://xenabrowser.net/datapages/). Patients with missing or insufficient data were excluded from this research. Finally, 178 tumor samples with immune cell infiltrating data were reserved for further analysis. To validate the established prognostic signature, the gene expression data normalized by RMA algorithm and clinical information of GSE21501 was obtained from GEO dataset (https://www.ncbi.nlm.nih.gov/geo/query/acc.cgi?acc=GSE57495) [12], a total 63 PAAD samples were included. The detailed clinic-pathological characteristics of PAAD patients from two datasets were exhibited in Tab. 1.

**Table 1.** Baseline characteristics of PAAD patients from TCGA and GSE57495 datasets.

| Characteristics | TCGA-PAAD |  | GSE57495 |  |
| --- | --- | --- | --- | --- |
|  | Cases | Proportion | Cases | Proportion |
| **Gender** |  |  |  |  |
| Female | 80 | 44.94% | 30 | 47.62% |
| Male | 98 | 55.06% | 33 | 52.38% |
| **Age** |  |  |  |  |
| ≤60 | 59 | 33.15% | - | - |
| >60 | 119 | 66.85% | - | - |
| **Subdivision** |  |  |  |  |
| Body | 14 | 7.87% | - | - |
| Head | 138 | 77.53% | - | - |
| Tail | 15 | 8.43% | - | - |
| Other | 11 | 6.18% | - | - |
| **Histology grade** |  |  |  |  |
| Well-Differentiated | 30 | 16.85% | 6 | 9.52% |
| Moderate-Differentiated | 96 | 53.93% | 35 | 55.56% |
| Poor-Differentiated | 50 | 28.09% | 18 | 28.57% |
| Unknown | 2 | 1.12% | 4 | 6.35% |
| **T stage** |  |  |  |  |
| T1 | 7 | 3.93% | 2 | 3.17% |
| T2 | 24 | 13.48% | 14 | 22.22% |
| T3 | 142 | 79.78% | 47 | 74.60% |
| T4 | 3 | 1.69% | - | - |
| Unknown | 2 | 1.12% | - | - |
| **N stage** |  |  |  |  |
| N0 | 50 | 28.09% | 30 | 47.62% |
| N1 | 123 | 69.10% | 32 | 50.79% |
| Unknown | 5 | 2.81% | 1 | 1.59% |
| **M stage** |  |  |  |  |
| M0 | 80 | 44.94% | - | - |
| M1 | 5 | 2.81% | - | - |
| Unknown | 93 | 52.25% | - | - |
| **TNM stage** |  |  |  |  |
| TNM 1 | 21 | 11.80% | 13 | 20.63% |
| TNM 2 | 146 | 82.02% | 50 | 79.37% |
| TNM 3 | 4 | 2.25% | - | - |
| TNM 4 | 5 | 2.81% | - | - |
| Unknown | 5 | 2.81% | - | - |
| **OS status** |  |  |  |  |
| Alive | 85 | 47.75% | 21 | 33.33% |
| Dead | 93 | 52.25% | 42 | 66.67% |

### 2.2. Immune infiltration analysis

ImmuCellAI (http://bioinfo.life.hust.edu.cn/web/ImmuCellAI/) is an emerging tool to estimate the abundance of 24 immune cells based on gene expression dataset [11]. Infiltrating data of TIICs corresponding to TCGA-PAAD samples was download from ImmuCellAI website. Besides, TIICs abundance of PAAD samples from GSE57495 was predicted by “Analysis” module of ImmuCellAI by uploading gene expression data.

### 2.3. LASSO Cox analysis

In survival analysis, overall survival (OS) events was set as end point of observation. To establish a TIICs-dependent prognostic signature, univariate Cox regression was first used to screen the prognostic values of 24 TIICs abundance. The least absolute shrinkage and selection operator (LASSO) Cox regression model was then applied for the further selection of prognostic TIICs. R package “glmnet” was used for LASSO analysis and establish the final model. Risk-score was calculated using a combination of the infiltrating abundance of TIICs and regression coefficients. Kaplan-Meier curves and log-rank analysis were used to identify survival differences by setting the cut-off value at median risk-score. Receiver operating characteristic (ROC) curves of the risk-score were generated using the R package “ROCR” to assess the prognostic accuracy of risk signature. Univariate and multivariable Cox analyses were used to study the independent prognostic value of risk-score combined with other clinic-pathological characteristics. At last, the GSE57495 data were used as the validation cohort to certify the effect of the risk-score signature.

### 2.4. Identification of differentially expressed genes

R language was applied to screen the DEGs between low-risk and high-risk samples using student t-test. The thresholds of extracting DEGs were as follows: |log2(fold change (FC))| >1, and P<0.05. Heatmap and volcano plot were drawn using the R package “heatmap” and “ggplot2”, respectively.

### 2.5. Function enrichment analysis

To get deep insight into the biological functions of critical genes, Gene ontology (GO) annotation analysis of DEGs was analyzed by the Database for Annotation, Visualization, and Integrated Discovery database (DAVID, http://david.ncifcrf.gov) [13]. The human genome (homo sapiens) was selected as the background variable. Enrichment terms were considered statistically significant when the P-value were less than or equal to 0.05. Top 20 terms was exhibited in Tab. 5.

**Table 2.** Univariate Cox regression analysis of TIICs infiltration in PAAD.

| **THCs** | **Beta** | **HR (95%CI for HR)** | **P-value** |
| --- | --- | --- | --- |
| CD4.naive | -320 | 1.5e-140 (0-Inf) | 0.750 |
| CD8.naive | -7 | 0.00089 (3.9e-08-20) | 0.170 |
| Tc | 3.3 | 26 (0.48-1400) | 0.110 |
| Tex | 11 | 70000 (0.65-7.6e+09) | 0.059 |
| Tr1 | -0.37 | 0.69 (0.015-32) | 0.850 |
| nTreg | 6.6 | 730 (1.4-380000) | **0.039** |
| iTreg | -0.71 | 0.49 (0.021-12) | 0.660 |
| Th1 | 4.5 | 88 (1.3-6100) | **0.038** |
| Th2 | -0.99 | 0.37 (0.00052-270) | 0.770 |
| Th17 | 5.6 | 280 (3.6-22000) | **0.011** |
| Tfh | -0.29 | 0.75 (0.13-4.2) | 0.740 |
| Tcm | 2.2 | 9.3 (0.14-630) | 0.300 |
| Tem | -8.9 | 0.00014 (3.7e-16-5.3e+07) | 0.510 |
| NKT | 1.1 | 3 (0.067-140) | 0.570 |
| MAIT | -4.1 | 0.017 (0.00045-0.64) | **0.028** |
| DC | 4.7 | 110 (3.2-3500) | **0.009** |
| B.cell | -1.2 | 0.31 (0.0067-14) | 0.550 |
| Monocyte | 5.3 | 210 (3.5-13000) | **0.011** |
| Macrophage | 1.8 | 5.9 (0.56-61) | 0.140 |
| NK | -0.057 | 0.94 (0.053-17) | 0.970 |
| Neutrophil | -1.6 | 0.2 (0.0031-13) | 0.450 |
| Tgd | -2.7 | 0.07 (0.00018-28) | 0.380 |
| CD4.T | -5.2 | 0.0056 (0.00013-0.24) | **0.007** |
| CD8.T | -4.5 | 0.011 (0.00014-0.87) | **0.043** |

**Table 3.**
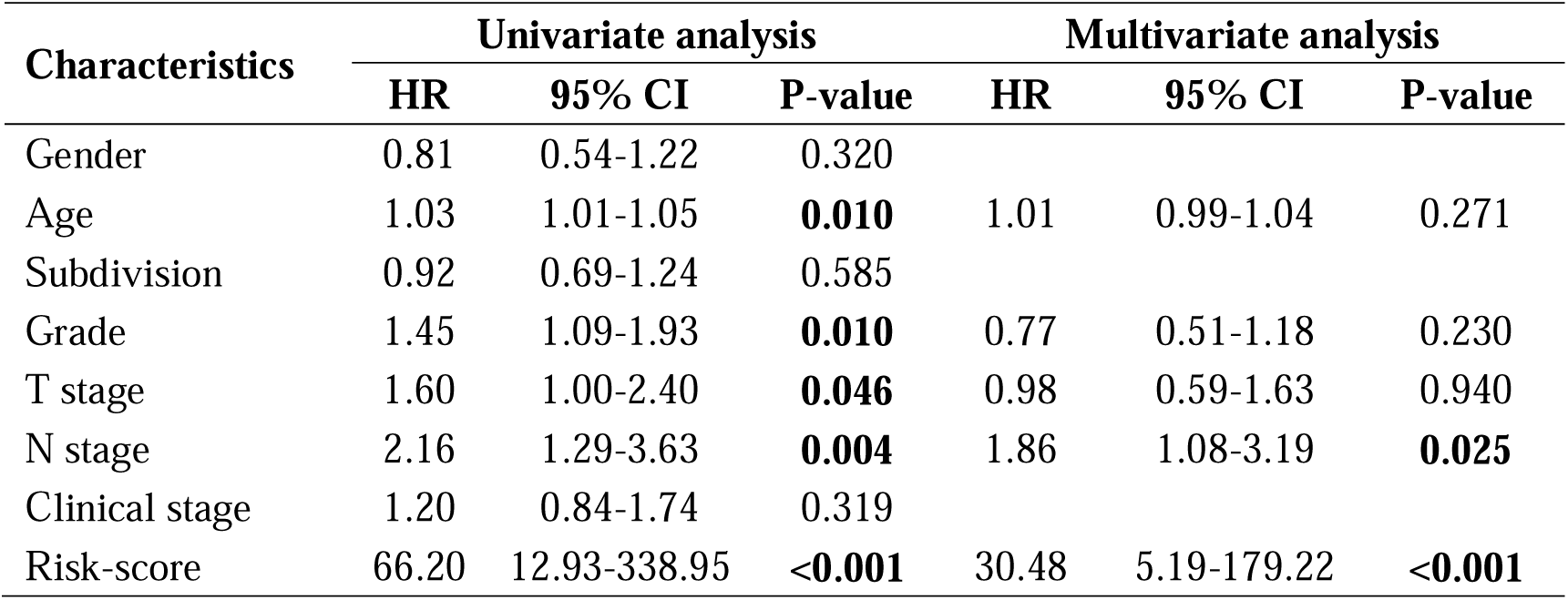
Univariate and multivariate analysis of survival factors in patients with PAAD.

| Characteristics | Univariate analysis |  |  | Multivariate analysis |  |  |
| --- | --- | --- | --- | --- | --- | --- |
|  | HR | 95% CI | P-value | HR | 95% CI | P-value |
| Gender | 0.81 | 0.54-1.22 | 0.320 |  |  |  |
| Age | 1.03 | 1.01-1.05 | **0.010** | 1.01 | 0.99-1.04 | 0.271 |
| Subdivision | 0.92 | 0.69-1.24 | 0.585 |  |  |  |
| Grade | 1.45 | 1.09-1.93 | **0.010** | 0.77 | 0.51-1.18 | 0.230 |
| T stage | 1.60 | 1.00-2.40 | **0.046** | 0.98 | 0.59-1.63 | 0.940 |
| N stage | 2.16 | 1.29-3.63 | **0.004** | 1.86 | 1.08-3.19 | **0.025** |
| Clinical stage | 1.20 | 0.84-1.74 | 0.319 |  |  |  |
| Risk-score | 66.20 | 12.93-338.95 | **<0.001** | 30.48 | 5.19-179.22 | **<0.001** |

**Table 4.** Association between risk-score and patients’ characteristics in PAAD.

| Characteristics | Cases | Low-risk | | High-risk | | $\chi^2$ | P-value |
| --- | --- | --- | --- | --- | --- | --- | --- |
|  |  | Cases | Proportion | Cases | Proportion |  |  |
| **Gender** |  |  |  |  |  | 0.363 | 0.547 |
| Female | 80 | 42 | 52.50% | 38 | 47.50% |  |  |
| Male | 98 | 47 | 47.96% | 51 | 52.04% |  |  |
| **Age** |  |  |  |  |  | 7.327 | **0.007** |
| ≤60 | 59 | 38 | 64.41% | 21 | 35.59% |  |  |
| >60 | 119 | 51 | 42.86% | 68 | 57.14% |  |  |
| **Subdivision** |  |  |  |  |  | 10.916 | **0.012** |
| Body | 14 | 6 | 42.86% | 8 | 57.14% |  |  |
| Head | 138 | 69 | 50.00% | 69 | 50.00% |  |  |
| Tail | 15 | 4 | 26.67% | 11 | 73.33% |  |  |
| Other | 11 | 10 | 90.91% | 1 | 9.09% |  |  |
| **Grade** |  |  |  |  |  | 3.553 | 0.314 |
| G1 | 30 | 18 | 60.00% | 12 | 40.00% |  |  |
| G2 | 96 | 49 | 51.04% | 47 | 48.96% |  |  |
| G3 | 48 | 22 | 45.83% | 26 | 54.17% |  |  |
| G4 | 2 | 0 | 0.00% | 2 | 100.00% |  |  |
| **T stage** |  |  |  |  |  | 5.635 | 0.131 |
| T1 | 7 | 4 | 57.14% | 3 | 42.86% |  |  |
| T2 | 24 | 17 | 70.83% | 7 | 29.17% |  |  |
| T3 | 142 | 65 | 45.77% | 77 | 54.23% |  |  |
| T4 | 3 | 1 | 33.33% | 2 | 66.67% |  |  |
| **N stage** |  |  |  |  |  | 1.673 | 0.196 |
| N0 | 50 | 21 | 42.00% | 29 | 58.00% |  |  |
| N1 | 123 | 65 | 52.85% | 58 | 47.15% |  |  |
| **TNM stage** |  |  |  |  |  | 5.221 | 0.156 |
| TNM1 | 21 | 14 | 66.67% | 7 | 33.33% |  |  |
| TNM2 | 146 | 71 | 48.63% | 75 | 51.37% |  |  |
| TNM3 | 4 | 1 | 25.00% | 3 | 75.00% |  |  |
| TNM4 | 5 | 1 | 20.00% | 4 | 80.00% |  |  |
| **OS status** |  |  |  |  |  | 14.073 | **<0.001** |
| Alive | 85 | 55 | 64.71% | 30 | 35.29% |  |  |
| Dead | 93 | 34 | 36.56% | 59 | 63.44% |  |  |

**Table 5.** Top 20 terms of BP enrichment of DEGs.

| **Category** | **Term** | **Count** | **P-value** |
| --- | --- | --- | --- |
| BP<sup>#</sup> | chemical synaptic transmission | 49 | **1.80E-12** |
| BP | cell adhesion | 72 | **5.70E-12** |
| BP | potassium ion transmembrane transport | 31 | **1.10E-10** |
| BP | regulation of insulin secretion | 20 | **2.80E-08** |
| BP | cell surface receptor signaling pathway | 44 | **5.90E-08** |
| BP | nervous system development | 45 | **8.40E-08** |
| BP | potassium ion transport | 21 | **1.90E-07** |
| BP | axonogenesis | 23 | **2.30E-07** |
| BP | synapse organization | 13 | **5.20E-07** |
| BP | regulation of ion transmembrane transport | 24 | **5.60E-07** |
| BP | axon guidance | 29 | **1.20E-06** |
| BP | chemokine-mediated signaling pathway | 18 | **2.10E-06** |
| BP | membrane depolarization during cardiac muscle<br>cell action potential | 8 | **5.70E-06** |
| BP | T cell costimulation | 18 | **8.10E-06** |
| BP | adaptive immune response | 26 | **9.30E-06** |
| BP | regulation of cardiac conduction | 15 | **9.50E-06** |
| BP | transmembrane receptor protein tyrosine kinase<br>signaling pathway | 20 | **1.10E-05** |
| BP | regulation of immune response | 29 | **1.10E-05** |
| BP | neuron migration | 21 | **1.20E-05** |
| BP | learning | 15 | **1.20E-05** |

#BP: Biological processes.

### 2.6. Protein-protein interaction network construction

The PPI network of DEGs was constructed using The Search Tool for the Retrieval of Interacting Genes (STRING) database (https://string-db.org/). The PPI network was visualized with Cytoscape software (version 3.7.2). The “Molecular Complex Detection” (MCODE) was applied to screen the key modules of the PPI network, and the top 2 modules was extracted for GO analysis.

### 2.7. Statistical analysis

R 3.6.3, SPSS 22.0, and GraphPad Prism 8.0 were applied as main tools for the statistical analysis and figures exhibition. The LASSO Cox regression model was employed for the further selection of prognostic TIICs by “glmnet”. Kaplan-Meier survival plots were generated with survival curves compared by log-rank test. The Chi-square test was used to evaluate differences in clinic-pathological variables between groups with different risks. Univariate and multivariate Cox regression models were used to calculate hazard ratio (HR) of risk-score and other clinic-pathological variables for OS. For DEGs screening, R language was applied using student t-test. For all analysis, differences were considered statistically significant when P-value was less than or equal 0.05.

## 3. Results

### 3.1. The distribution of TIICs in PAAD

To obtain a systematical insight into TIICs in HNSCC, the ImmuCellAI website was applied to calculate TIICs composition in PAAD simples in TCGA dataset. A heatmap was drawn to illustrate 24 immune cells proportions in these samples (Fig. 1A). As we see, the fraction of immune cells varied significantly among different samples. Due to the limited number of adjacent non-cancer samples, we failed to compare the difference of TIICs abundance in tumor and non-tumor samples. We next compared the proportion of different TIICs in PAAD samples. The results showed that type 1 regulatory T cell (Tr 1), natural killer cell (NK), macrophage, and *etc*. had higher abundance in PAAD, but CD4+ naïve cell, effector memory T cell (Tem), Tex cell, and *etc*. had lower abundance (Fig. 1B). Moreover, to evaluate the underlying relationships among different immune cell types, we assessed the correlation within TIICs in HNSCC. It was revealed that the fractions of several kinds of immune cells were mutually correlated in the TCGA-PAAD cohort (Fig. 1C). Overall, in view of different kinds of TIICs had distinct infiltrating features, we speculated that infiltrating TIICs might play various roles in regulating PAAD progression.

**Figure 1.**
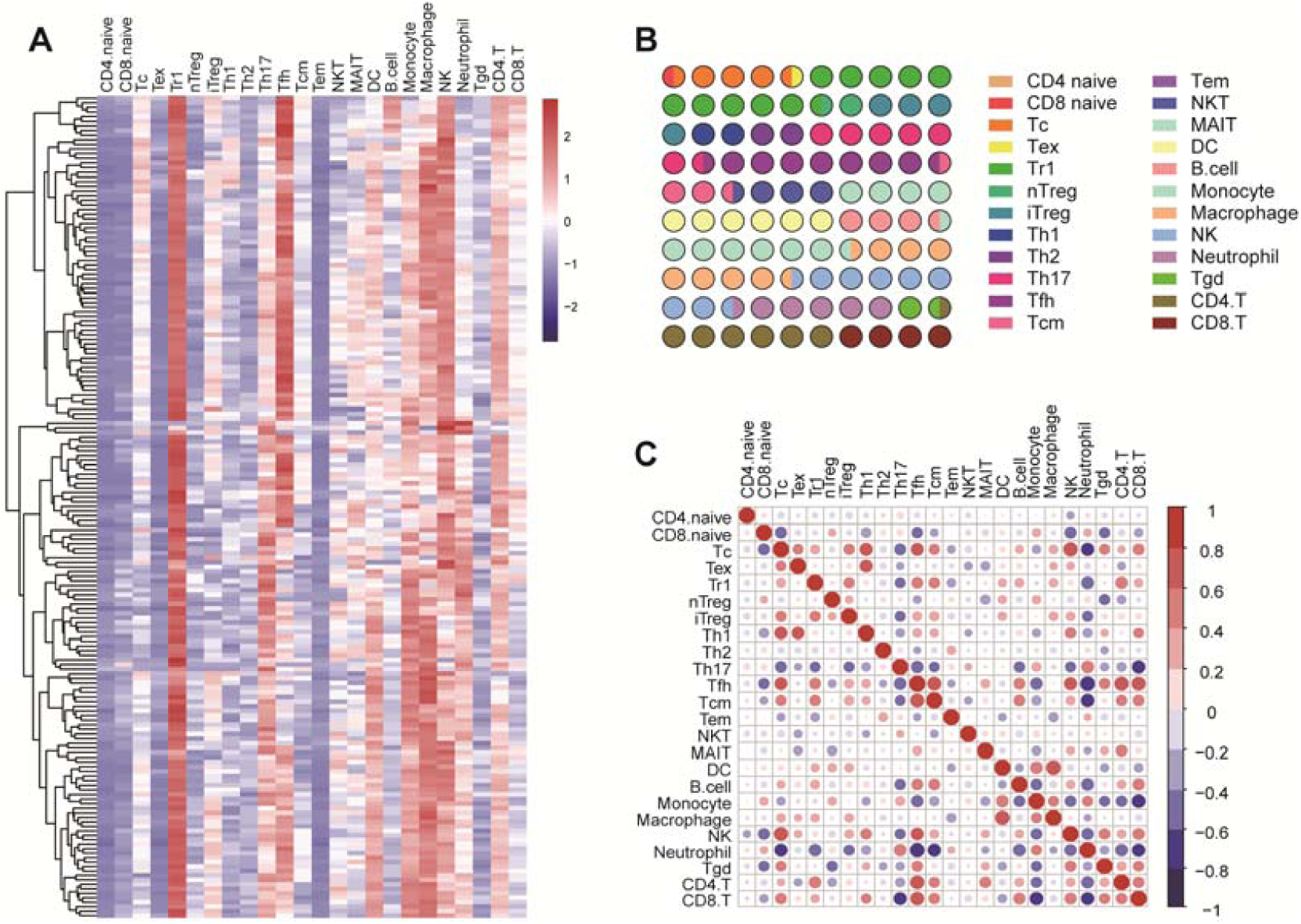
The distribution of TIICs in PAAD. (A) The heatmap of the 24 immune cells abundance based on TCGA-PAAD data. (B) The proportion of 24 immune cells in PAAD. (C) Correlation matrix of 24 immune cells abundance in PAAD samples.

### 3.2. The prognostic values of TIICs in PAAD

To assess the prognostic values of infiltrating TIICs for PAAD patients, survival analysis with log-rank test was applied based on TCGA-PAAD data. The patients were divided into two groups according to the median infiltration levels of TIICs. The result showed that most TIICs had no obvious prognostic values in PAAD patients (Fig. 2A). However, patients with higher levels of Tex and monocyte had significantly worse OS (Figs. 2B, 2D), while high infiltrating levels of mucosal-associated invariant T cell (MAIT) and CD4+ T cell predicted better prognosis in PAAD patients (Figs. 2C, 2E). Taken together, these findings suggested that several TIICs had specific prognostic values in PAAD.

**Figure 2.**
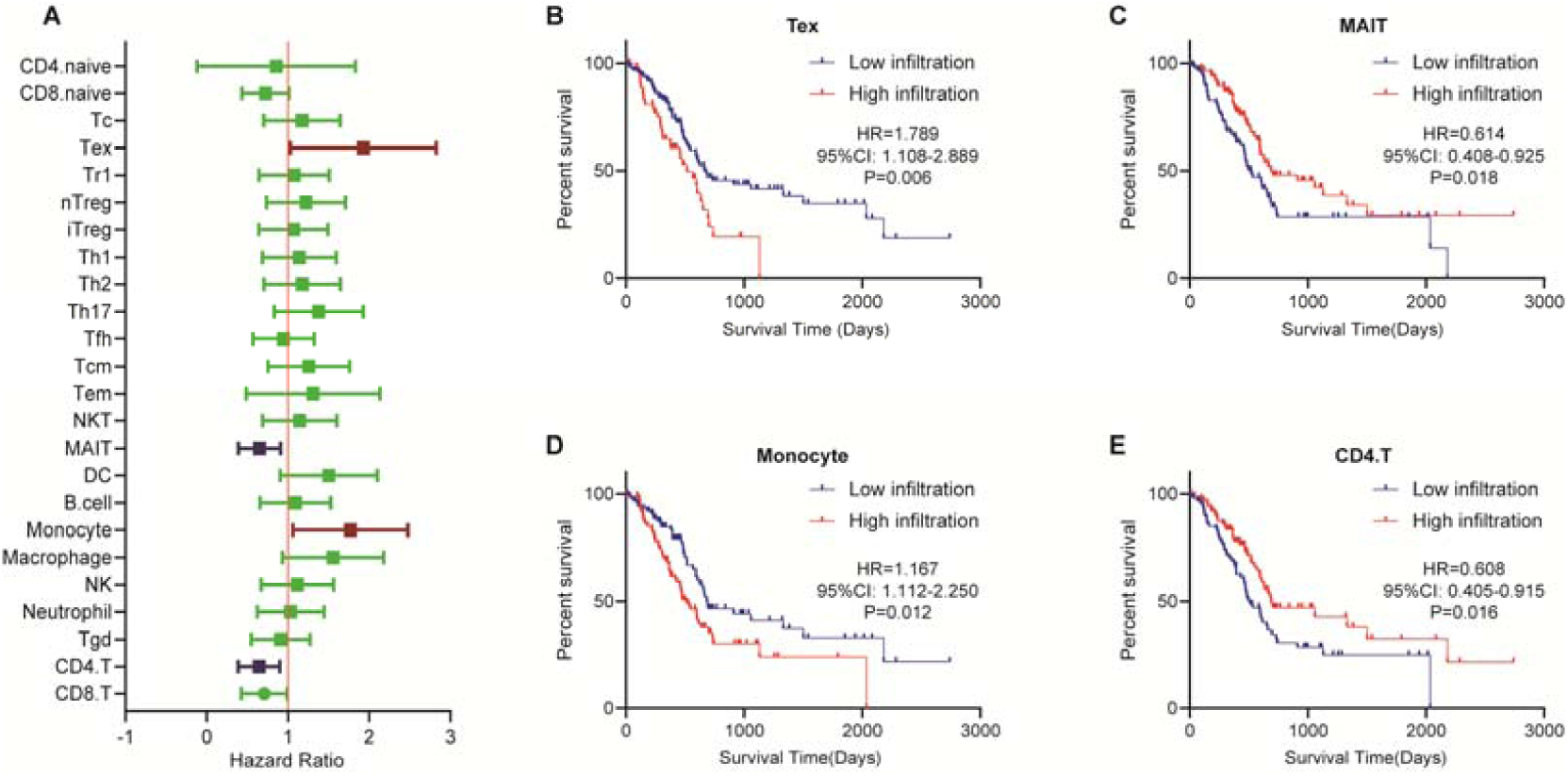
Survival plots of TIICs in PAAD patients. (A) Overview of Kaplan-Meier analysis for the prognostic values of 24 selected TIICs in PAAD. (B) High Tex infiltration predicted worse prognosis in PAAD patients. (C) Low MAIT infiltration predicted worse prognosis in PAAD patients. (D) High monocyte infiltration predicted worse prognosis in PAAD patients. (E) Low CD4+ T cell infiltration predicted worse prognosis in PAAD patients.

### 3.3. Establishment and validation of a TIICs signature

In view of the prognostic values of TIICs, we next try to establish a TIICs associated prognostic signature. We first conducted univariate Cox regression to initially screen TIICs with the significant impact on PAAD prognosis. A total 8 TIICs had promising prognostic impact in PAAD (Tab. 2). Subsequently, LASSO Cox analysis with ten-fold cross-validation was performed in the TCGA-PAAD dataset to further narrow the effective TIICs (Fig. 3A, 3B). Six TIICs were identified and subsequently used to construct a prognostic signature. We next constructed a 6-TIICs signature to assess the prognosis of PAAD patients based on the infiltrating levels of these 6 TIICs and their regression coefficients (Fig. 3C).

**Figure 3.**
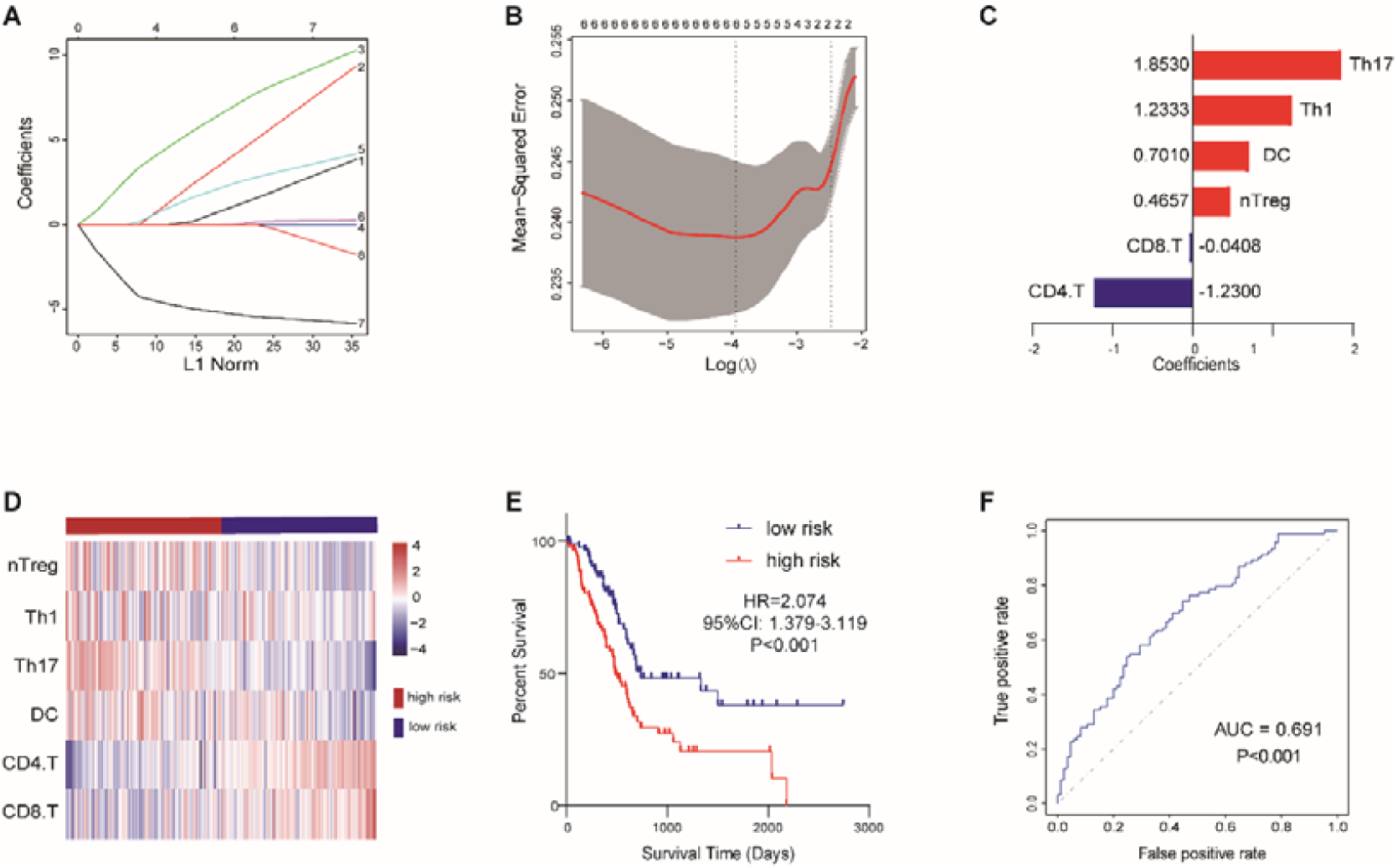
The establishment of TIICs-related signature in PAAD. (A) LASSO coefficient profiles of 8 selected TIICs. (B) 10-fold cross-validations result which identified optimal values of the penalty parameter λ. (C) The distribution of LASSO Cox coefficients in the TIICs-related signature. (D) The heatmap of six TIICs infiltrating levels in the training set. (E) Patients in the high-risk group exhibited worse OS compared to those in the low-risk group in the training set. (F) ROC analysis in the training set.

Patients in the TCGA cohort were divided into low-risk group (n=89) and high-risk group (n=89) utilizing the median risk-score as the cut-off value. Infiltrating abundance of these six TIICs had obvious distinction between two groups (Fig. 3D). The Kaplan-Meier curves exhibited that high-risk patients had notably poor prognosis in the training set (P<0.001, Fig. 3E). The multivariate Cox analysis uncovered that this prognostic signature was an independent prognostic factor for PAAD patients (Tab. 3). We next conducted ROC analysis to assess the prognostic accuracy of the risk-score, and the result showed that the risk-score had a nice prognostic accuracy (Fig. 3F). Moreover, Chi-square test showed that the risk-score was associated with several clinic-pathological features, such as age, tumor subdivision and OS status (Tab. 4).

To further validate the prognostic value of signature in PAAD, we next used 63 patients from GSE57495 dataset as the validation cohort. Similar to training cohort, the infiltrating abundance of these six TIICs had obvious distinction between low-risk group (n=32) and high-risk group (n=31) (Fig. 4A). Kaplan-Meier curves suggested that the high-risk group patients had significantly worse outcomes than low-risk group (Fig. 4B), and the ROC analysis verified that the risk-score had a good prognostic accuracy (Fig. 4C). To sum up, our results indicated a satisfactory value of the TIICs-associated signature for survival prediction.

**Figure 4.**
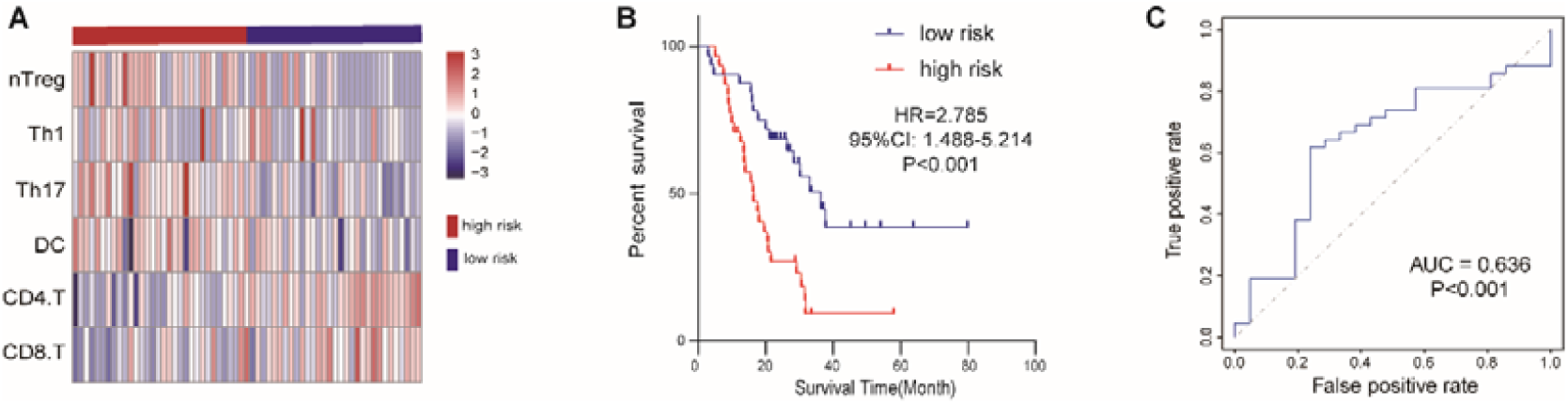
Validation of the signature in GSE57495 cohort. (A) The heatmap of six TIICs infiltrating levels in the validated set. (B) Patients in the high-risk group exhibited worse OS compared to those in the low-risk group in the validated set. (C) ROC analysis in the validated set.

### 3.4. Identification of DEGs and enrichment analysis

To further understand the progression of PAAD, we exacted DEGs between two groups. Base on the stated threshold, a total 1,334 DEGs were identified, including 1,082 up-regulated genes and 252 down-regulated genes, as shown in heatmap and volcano plot (Figs. 5A, 5B). Then, the GO enrichment analysis was performed for the 1,334 DEGs. The biological processes mainly involved in chemical synaptic transmission, cell adhesion, potassium ion transmembrane transport, regulation of insulin secretion and so on (Fig. 5C, Tab. 5). Besides, several immune-related biological processes were also regulated by DEGs, such as chemokine-mediated signaling pathway, T cell costimulation, and adaptive immune response (Tab. 5).

**Figure 5.**
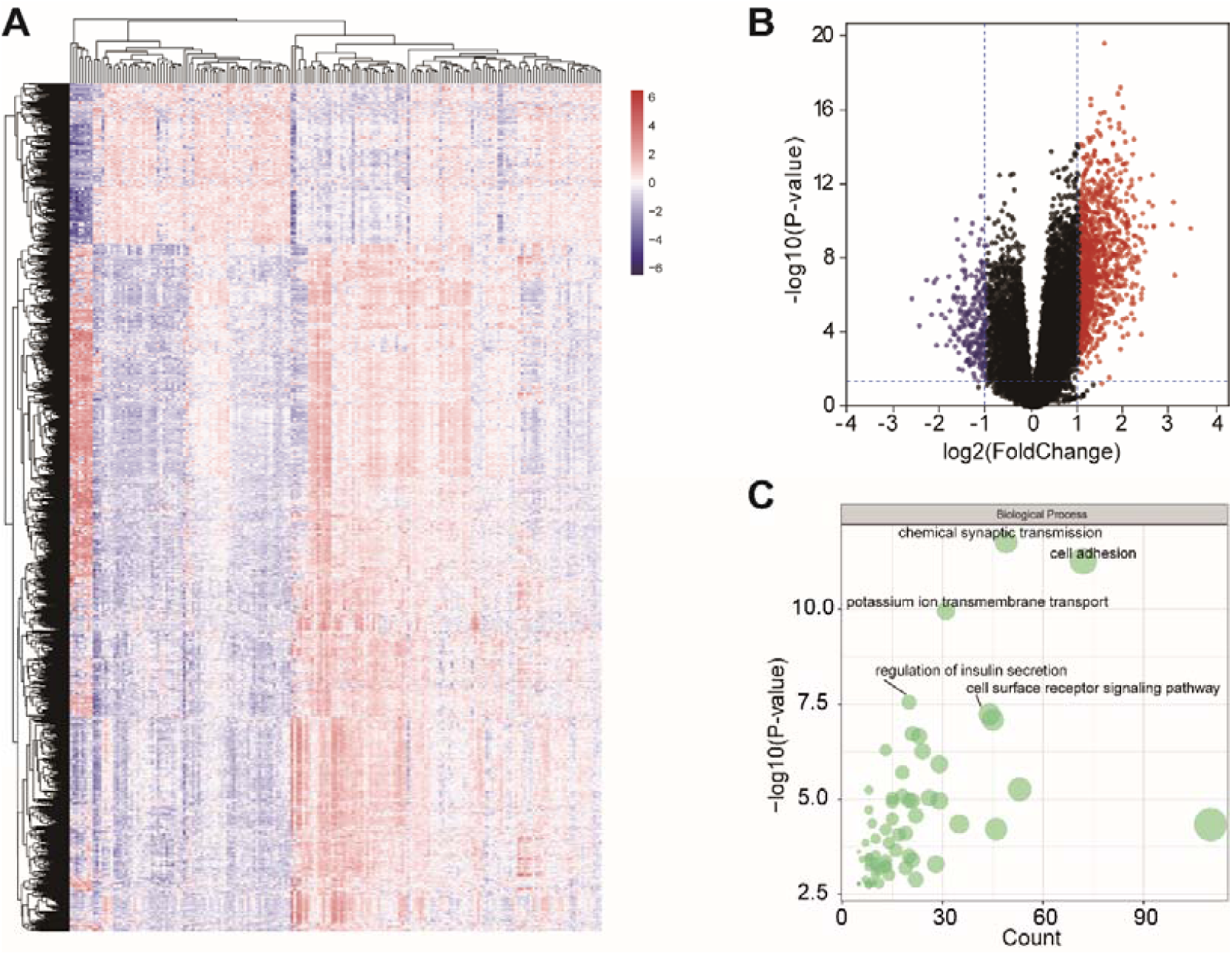
DEGs between low- and high-risk groups. (A) The heatmap of DEGs in PAAD. (B) The volcano plot of DEGs with the thresholds of |log2(fold change (FC))| >1 and P < 0.05. (C) BP enrichment analysis of total DEGs.

### 3.5. PPI construction and biological function of critical modules

The PPI network of 1,334 DEGs was constructed by STRING database to predict the potential interactions of DEGs. To obtain hub genes that had tightest interactions with each other, we conducted the MCODE analysis to identify the top two modules in the PPI network (Figs. 6A, 6B). The main BP terms analysis of DEGs in top one module were mainly involved in G-protein coupled receptor signaling pathway, chemokine-mediated signaling pathway, chemotaxis, inflammatory response, and immune response (Fig. 6A). The other hub module were mainly associated with regulation of immune response, cell surface receptor signaling pathway, T cell costimulation, T cell activation, and adaptive immune response (Fig. 6B). Taken together, these findings provided a novel understand about the progression of PAAD from the insight of immune-related biological processes.

**Figure 6.**
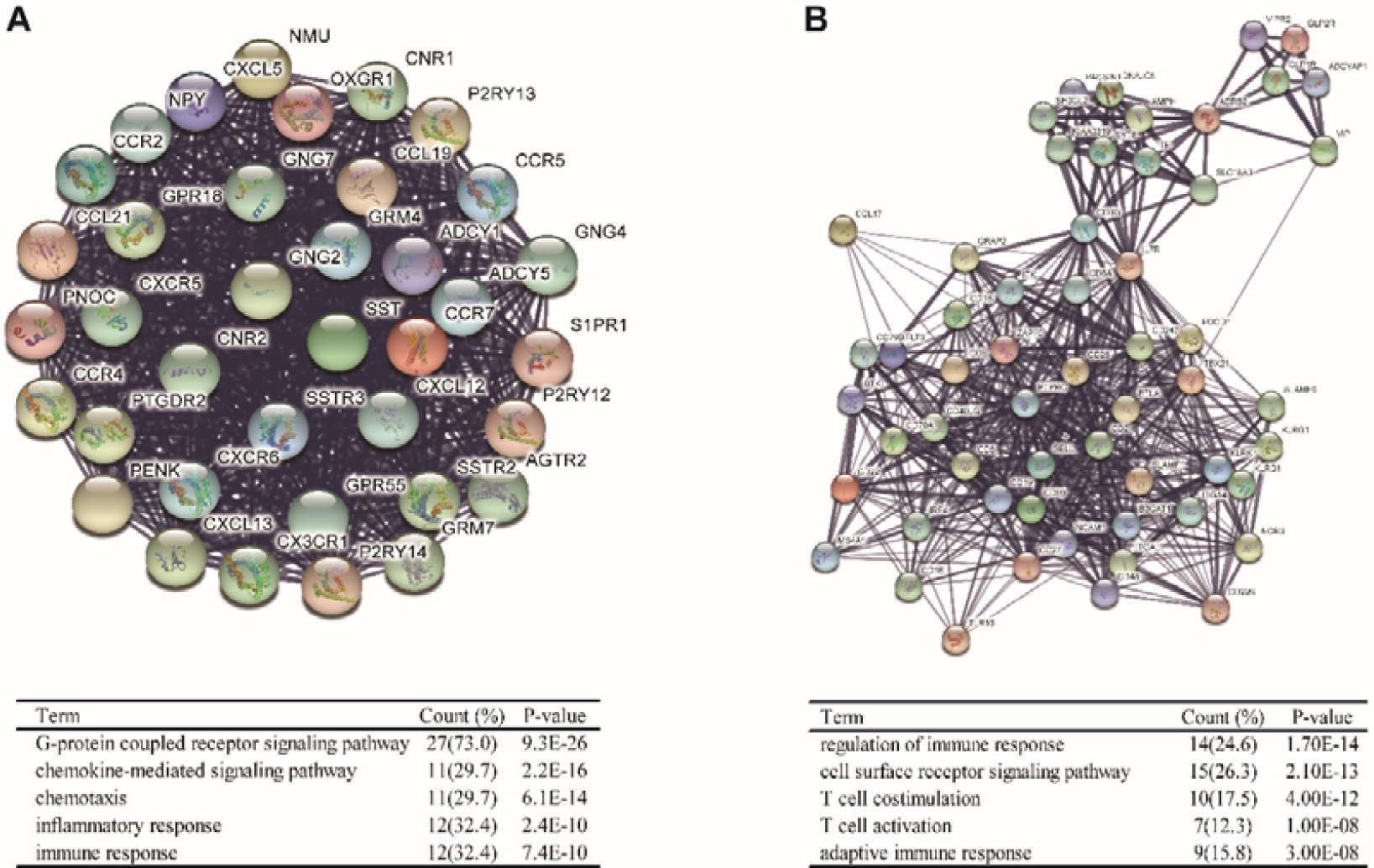
Two key modules of PPI network and related biological functions. (A) Module 1 (nodes: 37, score: 37.000), (B) Module 2 (nodes: 57, score: 20.821). Note: Top 5 biological processes terms were exhibited in this Figure.

## 4. Discussion

As well all known, immune cells infiltrating in the tumor microenvironment have been uncovered to play prominent roles in the biological behaviors of various cancers [14-16]. It has been revealed that identifying subtypes of the immune microenvironment in PAAD provides the promising opportunities for therapeutic development based on personalization of systemic immunotherapies [17]. Thus, summarizing the features of immune cells infiltrating in tumor microenvironment is essential for risk classification, treatment, and prognosis assessment for PAAD. In the current study, we assessed infiltrating levels of 24 immune cells and explored their potential prognostic values in PAAD through the ImmuCellAI algorithm.

The TIICs are mainly composed of B cell, T cell, macrophage, monocyte, NK and *etc*., which act as significant roles in promoting and/or suppressing cancer progression [18, 19]. B cells, producers of antibody, not only are significant components of the adaptive immune system, but also could co-operate with other TIICs by secreting cytokine and presenting antigen [20]. T cells contain a large family, including CD8+ T cell, CD4+ T cell, Th, Treg and MAIT. Most T cell subsets play critical roles in tumor control [21], but several kinds of T cells can also promote the cancer progression. For example, as a member of CD8+ T cell subsets, Tex lose robust effector functions by expressing multiple inhibitors, which are defined by alterant transcriptional programs [22]. Previous study uncovered that high level of CD8+ Tex expressing PD1 predicted worse clinical outcome in hepatocellular carcinoma [23]. A mass of studies have demonstrated that infiltrating abundance of specific TIICs are associated with improved or unfavorable clinical outcomes in different cancers [24-26]. In this study, we found that high infiltrating levels of Tex and monocyte predicted worse OS, while low abundance of MAIT and CD4+ T cell were associated poor prognosis in PAAD patients, suggesting several types of TIICs had encouraging prognostic impacts on PAAD.

As well all known, PAAD has been identified as a type of aggressive malignancy with awfully poor prognosis. Growing numbers of studies attempt to systematically summarize the malignant characterization of PAAD and develop prognostic risk identifiers from multi-omics perspective [27, 28]. Besides, immune-related genes are also treated as hotspots in the field of prognostic assessment. Zhang *et al*. developed a PD-L2-based immune signature to exactly predict survival in resected PAAD [29]. Meng *et al*. screened DEGs between ESTIMATE score groups, and further established an 8-mRNA signature prognostic identifier for PAAD [30]. However, as far as we know, there is no exploration on prognostic values of combinations of multiple immune cells in cancers up to data. In this report, we developed a TIICs-related prognostic signature, consisting of Th17, Th1, DC, nTreg, CD8+ T cell, and CD4+ T cell, which could precisely indicated prognosis in PAAD patients. These findings suggested that different TIICs combinations might obtain better predictive values in prognostic assessment for cancerous diseases.

Additionally, we screened the risk-related DEGs and found that biological processes of these DEGs biological processes mainly involved in chemical synaptic transmission, cell adhesion, potassium ion transmembrane transport, regulation of insulin secretion, and so on. Then, we exacted the hub gene modules by Cytoscape software, and found that the top 2 modules were significantly associated with immune response. It has been confirmed that PAAD microenvironment was highly immunosuppressive point and inhibited immune escape might be critical to control PAAD progression [31]. In this study, we explored the biological functions of DEGs between different risk groups, which might explain the possible mechanisms of malignant progression of PAAD from the view of immune-related factors.

To sum up, we analyzed the 24 TIIC subgroups in PAAD samples based on transcriptome data using ImmuCellAI tool. As an important result, we identified and validated a six-TIICs prognostic signature, which could precisely predict prognosis in PAAD patients. However, although algorithms based on transcriptional data may be able to predict the abundance of TIICs with great accuracy, large-scale clinical investigation by single cell sequencing and/or multicolor immunofluorescence technologies should be conducted to further validate the findings of the current research.

## Abbreviation

PAAD: pancreatic cancer;
TIICs: tumor-infiltrating immune cells;
RNA-Seq: RNA sequencing;
Treg: regulatory T cell;
Tc: cytotoxic T cell;
Tex: exhausted T cell;
Th1: T helper 1 cell;
DC: dendritic cell;
DEGs: differentially expressed genes;
PPI: protein-protein interaction;
OS: overall survival;
LASSO: least absolute shrinkage and selection operator;
ROC: Receiver operating characteristic;
FC: fold change;
GO: gene ontology;
DAVID: Database for Annotation, Visualization, and Integrated Discovery;
STRING: Search Tool for the Retrieval of Interacting Genes;
MCODE: Molecular Complex Detection;
HR: hazard ratio;
Tr 1: type 1 regulatory T cell;
NK: natural killer cell;
Tem: effector memory T cell;
MAIT: mucosal-associated invariant T cell.;

## Acknowledgments

This work was founded by the Natural Science Foundation of Jiangsu Province of China (BE2017626), the Six Talent Peaks Project of Jiangsu Province (WSN-186), and the Foundation of Wuxi Health Commission (QNRC003).

